# Knowledge and attitudes of Ebola among the general public of Trinidad and Tobago during the 2014-15 West Africa outbreak

**DOI:** 10.1101/346999

**Authors:** S Pooransingh, S Mohammed, K Melville, C Mohammed, M Mohammed, R Mohammed, W Mootoo, M Morris, D Motilal, B Bhagwandeen, I Dialsingh

## Abstract

**Objective:** Health system resilience and resilience of a country include the capacity of health personnel, institutions, and populations to prepare for and effectively respond to crises. This study investigates the knowledge and attitudes of the public concerning Ebola Virus Disease in Trinidad and Tobago.

**Design and Methods:** A cross sectional study whereby respondents (n = 920) were sampled from public places. Data were collected via interviewer administered questionnaires. Data were analysed using SPSS version 23.

**Results:** The response rate was 67.6 % (622/920). The main age category of responders was the 20 to 30 year age category (40.5%); responders were mostly female (58.0 %). Regarding knowledge, there were significant differences among occupational categories (F = 2.811, df1 = 6, df2 = 571, p-value = 0.011). Tukey’s HSD post hoc test revealed that the mean knowledge scores for professional and sales occupations differed significantly (p-value = 0.003). There was a significant association between being afraid to go for treatment and age (p-value = 0.001). Significant associations were also found between occupational grouping and education attainment with opinion about the preparedness of private medical facilities, likelihood to shun family members with Ebola, being afraid to go for treatment and preference for traditional medicine (p-value <0.05).

**Conclusion:** This study highlights opportunities for community engagement to enhance health system resilience during outbreaks which would maximise national and global health security.

## Introduction

Countries have been working to strengthen their alert and response capacities for acute public health threats since 2007 under the International Health Regulations (IHR 2005) (1). Several events have served to test these efforts such as the H1N1 Pandemic 2009, the 2011 Japan earthquake and tsunami, the emergence of Middle East Respiratory Syndrome (MERS) in 2012, various global terrorist events, Zika in the Americas in 2015 and the Ebola (EVD) outbreak in West Africa in 2014/5 (1). Ebola constituted a Public Health Emergency of International Concern as it left Africa for the first time, presenting a real threat to other countries. No longer could Ebola be considered a disease of ‘over there’ (2). The Ebola epidemic raised to the fore the fragility of health systems. It showed what happens when a crisis hits and the health systems are not prepared (3).

EVD is highly contagious with human to human spread via direct bodily contact with the blood, secretions, organs or other bodily fluids of infected people, and with surfaces and materials (e.g. bedding, clothing) contaminated with these fluids. The case fatality rate is high if persons are not treated (1,4).

With increased air travel and persons from West Africa setting abroad including in Trinidad and with links to their home country, the likelihood of a case or a contact arriving into the country or any country, though rare, is now possible. Countries need to increase their resilience i.e. the intrinsic capacity of individuals, populations and infrastructure to rebound from shocks (5,6).

The status of Trinidad and Tobago’s health systems preparedness is known from annual IHR 2005 status reports; it was therefore decided to seek the public’s knowledge and attitudes on Ebola as the community is a central player in in the emergency response and a successful response relies on the participation and cooperation of the public. A well informed and well-rehearsed public would be better able to act faster and effectively.

## Materials and Methods

### Study Design

A cross-sectional survey was undertaken.

### Study Period

The survey was undertaken from January to May 2016.

### Study setting

The study was undertaken in public spaces in Trinidad and Tobago (TT), a twin island republic in the chain of islands collectively termed the Caribbean. The population of TT is approximately 1. 4 million persons (7).

### Study Population

The study population comprised the adult (18 years and older) population of Trinidad and Tobago. This included males and females of varying ethnic compositions, representative of the general population. Responders were chosen by convenience sampling. Questionnaires were distributed along busy streets, at malls, near bus stops, as well as other high traffic areas across the different regions of Trinidad and Tobago.

### Sample size

The sample size for the study was initially calculated at the onset as 913 responders.

### Data Collection

A total of 920 questionnaires were distributed across Trinidad and Tobago. Responders were briefed on the study’s objectives before verbal consent was obtained for participation. Interviewers assured the responders that their responses would be anonymous, that there would be no way identifying them through the results. The questionnaire collected demographic information, general knowledge and attitudes towards Ebola and views of responders on the healthcare system in the country.

### Data analysis

All data collected were analyzed using SPSS version 23. A knowledge scale was constructed using the sum total score of nineteen binary questions concerning respondents’ knowledge about EVD. The two-sample independent t-test was used to compare the mean knowledge score between males and females. An ANOVA test was used to determine whether there were differences among categories in the age, occupation and education variables. Tukey’s HSD Procedure was carried out to determine which pairs of mean scores across occupational categories differed (adjusted p-value <0.05). In order to determine if there was a relationship between the attitudes toward Ebola and demographic factors (age, gender, occupation and education), Chi-Squared Tests of Independence were carried out. In the cases where at least 20% of the cells had expected counts less than 5, Fisher’s Exact Test was used.

### Ethical Approval

The University of the West Indies, St. Augustine, Trinidad and Tobago Ethics Committee granted ethical approval for this study (Reference: CEC063/11/15).

## FINDINGS

A total of 920 participants were approached to participate of which 622 (67.6%) agreed. There were 42% male and 58% female responders. The ethnic groups of the responders comprised those of East Indian origin (35.9%), African origin (27.2%) and other, including mixed ethnicity (36.9%). The majority of responders reported attaining tertiary education (55.6%) followed by secondary education (41.1%). Table 1 summarises the age breakdown of responders and Table 2 the occupational categories.

**Table 1.** Table showing the age and occupational groupings breakdown of responders.

| Age Group (years) | Frequency (%) | Occupational Category | Frequency (%) |
| --- | --- | --- | --- |
| <20 | 74 (11.2) | Managers/Directors/Senior Officials | 58 (9.9) |
| 20 – 30 | 267 (40.5) | Professional Occupations | 78 (13.3) |
| 31 – 40 | 128 (19.4) | Administrative/Secretarial | 32 (5.5) |
| 41 – 50 | 87 (13.2) | Skilled Trades | 17 (2.9) |
| >50 | 104 (15.7) | Caring/Leisure/Other | 293 (50.1) |
|  |  | Sales/Customer Service | 102 (17.4) |
|  |  | Elementary Occupations | 5 (0.9) |
| **Total** | **660 (100.0)** |  | **585 (100.0)** |

### Findings related to Knowledge about Ebola

In terms of knowledge questions about Ebola, more than 70% of responders knew what EVD was and how it is transmitted. Forty-five per cent (45%) of responders reported that there is a cure for EVD. The average knowledge score, based on 622 (67.6%) participants, was 10.61 (sd = 3.756). A Cronbach’s alpha coefficient of 0.681 indicated the constructed knowledge scale was a relatively consistent measure of the concept.

There were no differences in the knowledge scores between males and females (T=−0.959, df= 608, p-value = 0.338).

There were no significant differences in the knowledge scores among the five age groups (F = 2.186, df1 = 4, df2= 613, p-value = 0.069). However, there were significant differences among the mean knowledge score of the seven occupational categories (F = 2.811, df1 = 6, df2 = 571, p-value = 0.011). It was found that the mean knowledge scores for professional and sales occupations differed significantly (Tukey’s HSD adjusted p-value = 0.003). There were also significant differences among education categories (p value = <0.001). Table 2 summarizes the results.

**Table 2.**
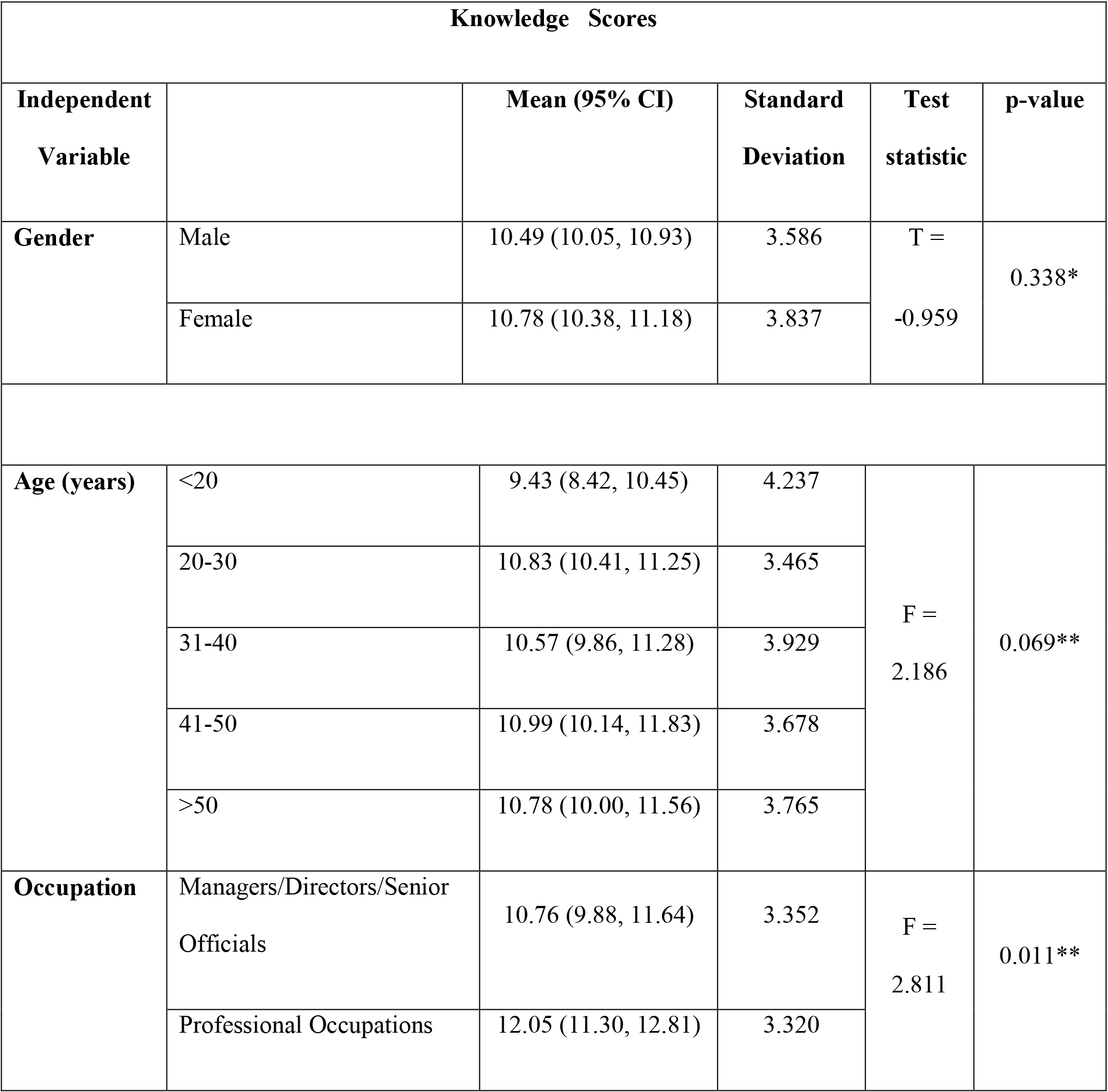
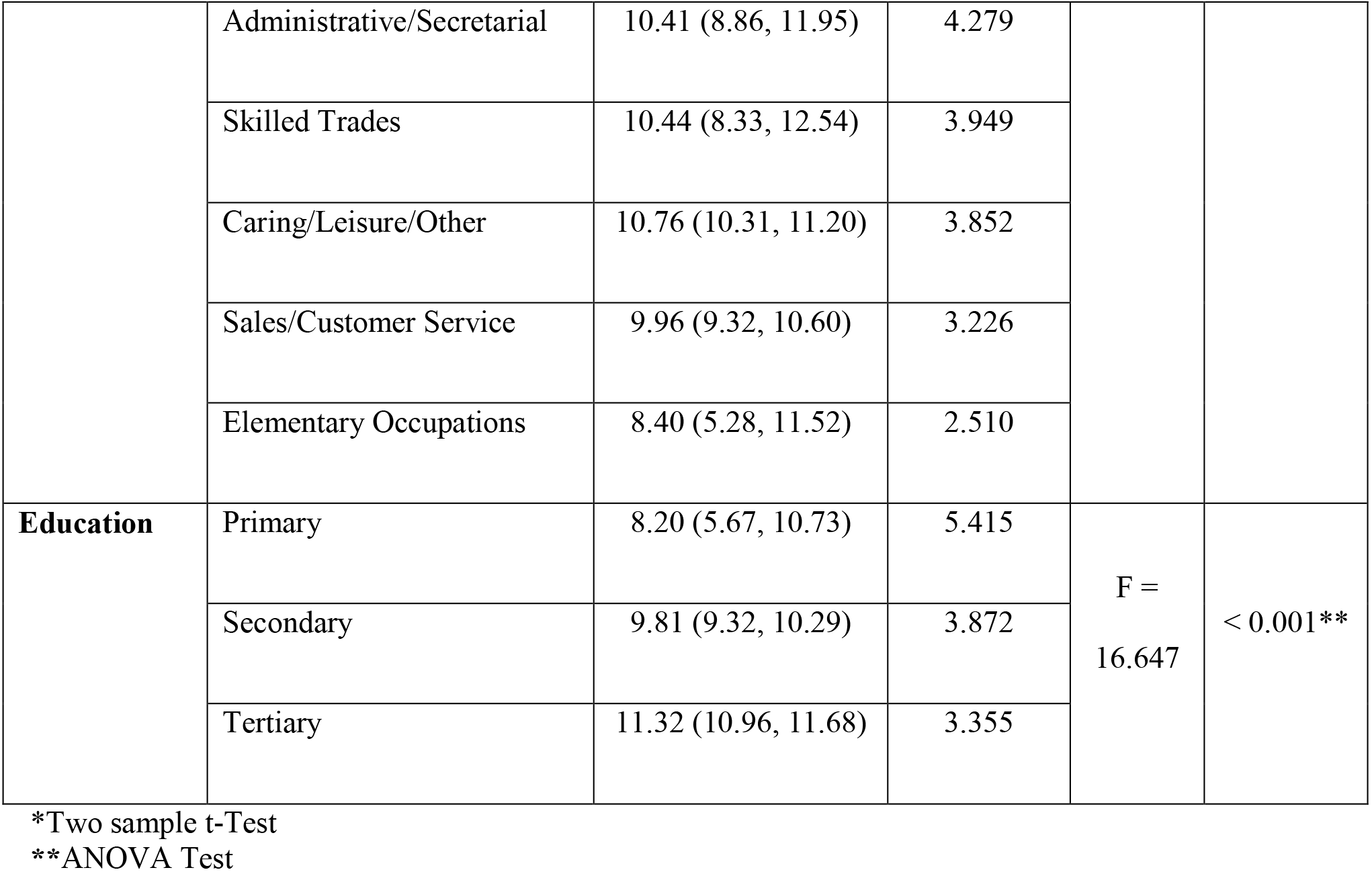
Knowledge of Ebola Scores versus Age, Gender, Occupation and Education.

### Findings related to Attitudes towards Ebola

Of the 586 participants, 50% believed that an outbreak of the EVD was very likely in TT and the most common reason cited for this was easy travel between countries. Furthermore, 97% thought the citizens of TT were not prepared for an outbreak. In terms of age groups, 355 (60.1%) of the 591 respondents were between the ages of 20 and 40 years and believed the country is not prepared to deal with an outbreak of the EVD. It was found that opinions on the likelihood of an outbreak were not dependent on gender, age, occupation and education (p-values > 0.05).

The top three reasons given for the country being ill-prepared for an outbreak were outdated healthcare facilities, lack of education and awareness, and ignorance and laziness among the public.

The majority of participants thought both private and public health care facilities were not prepared to deal with an outbreak. They reported that an outbreak would result in death and panic.

The majority of responders (62%) would be likely or very likely to shun someone with the disease; while 21% were neutral in response. When asked about a family member with EVD, 24% were neutral, and 35% likely or very likely to shun their family member.

Almost one fifth of responders said they would be afraid to go for treatment; 14% stated they would opt for traditional medicine over a health facility. There was no association between citizens who believed there is a cure for the disease and those who would rather use traditional medicine than go to a medical facility (p-value = 0.307).

Being afraid to go for treatment was significantly associated with age of the respondent (p-value = 0.001).

Occupational category was found to be significantly associated with opinion towards private medical facilities preparedness, likelihood to shun family members, being afraid to go for treatment and preference for traditional medicine (p-value < 0.05).

Education was found to be significantly associated with opinion towards private medical facilities preparedness, likelihood to shun a person with EVD, being afraid to go for treatment and preference for traditional medicine (p value < 0.05).

No association with any aspect of attitude was found with gender (p-values > 0.05).

**Table 3.** Association of Attitudes toward Ebola and Age Category.

| Variable |  | Age (years) |  |  |  |  | p-value |
| --- | --- | --- | --- | --- | --- | --- | --- |
|  |  | <sup>1</sup><20 | 20-30 | 31-40 | 41-50 | >50 |  |
| Public medical facilities prepared? | No | 63 | 242 | 107 | 71 | 78 | 0.926*** |
|  | Yes | 2 | 9 | 6 | 3 | 4 |  |
|  | **Odds Ratio (95% C.I.)** |  | 1.171 (0.247, 5.558) | 1.766 (0.346, 9.018) | 1.331 (0.215, 8.223) | 1.615 (0.287, 9.108) |  |
| Private medical facilities prepared? | No | 46 | 194 | 86 | 62 | 69 | 0.477**** |
|  | Yes | 15 | 48 | 19 | 9 | 14 |  |
|  | **Odds Ratio**<br>**(95% C.I.)** |  | 0.759 (0.391,<br>1.472) | 0.678<br>(0.315,<br>1.457) | 0.445<br>(0.179,<br>1.106) | 0.622 (0.275,<br>1.410) |  |
| **Likely to shun**<br>**person?** | Not Very Likely | 3 | 24 | 12 | 8 | 9 | 0.414**** |
|  | Not Likely | 5 | 20 | 8 | 2 | 3 |  |
|  | Neutral | 19 | 55 | 19 | 9 | 13 |  |
|  | Likely | 7 | 36 | 16 | 9 | 13 |  |
|  | Very Likely | 24 | 106 | 51 | 39 | 42 |  |
| **Likely to shun**<br>**family**<br>**member?** | Not Very Likely | 11 | 53 | 29 | 16 | 22 | 0.775**** |
|  | Not Likely | 10 | 47 | 28 | 12 | 18 |  |
|  | Neutral | 17 | 58 | 25 | 17 | 25 |  |
|  | Likely | 12 | 39 | 16 | 12 | 10 |  |
|  | Very Likely | 17 | 55 | 18 | 12 | 12 |  |
| **Afraid to go**<br>**for treatment?** | Strongly<br>Disagree | 32 | 136 | 72 | 41 | 62 | 0.001**** |
|  | Disagree | 15 | 46 | 18 | 11 | 6 |  |
|  | Neutral | 12 | 26 | 8 | 1 | 4 |  |
|  | Agree | 1 | 22 | 6 | 4 | 1 |  |
|  | Strongly Agree | 7 | 24 | 12 | 12 | 15 |  |
| **Rather use**<br>**traditional**<br>**medicine?** | Strongly<br>Disagree | 33 | 138 | 68 | 42 | 62 | 0.402**** |
|  | Disagree | 11 | 35 | 19 | 10 | 8 |  |
|  | Neutral | 11 | 44 | 13 | 7 | 8 |  |
|  | Agree | 5 | 18 | 9 | 5 | 1 |  |
|  | Strongly Agree | 5 | 19 | 8 | 5 | 9 |  |

\*\*\*Fisher's Exact Test
\*\*\*\* $\chi^2$ Test
<sup>1</sup>Reference Category

**Table 4.**
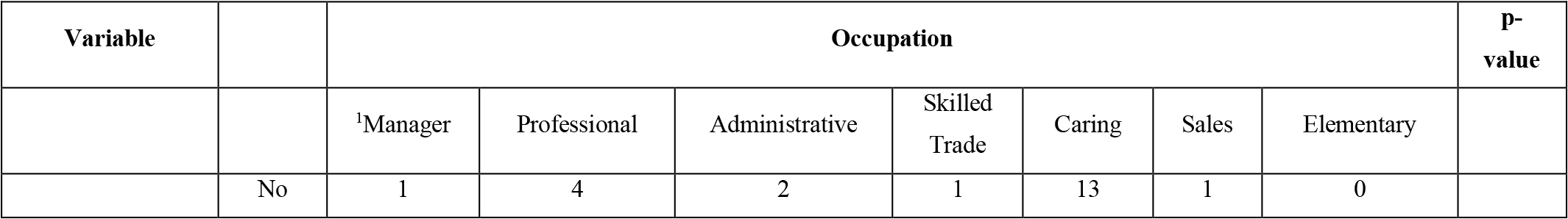
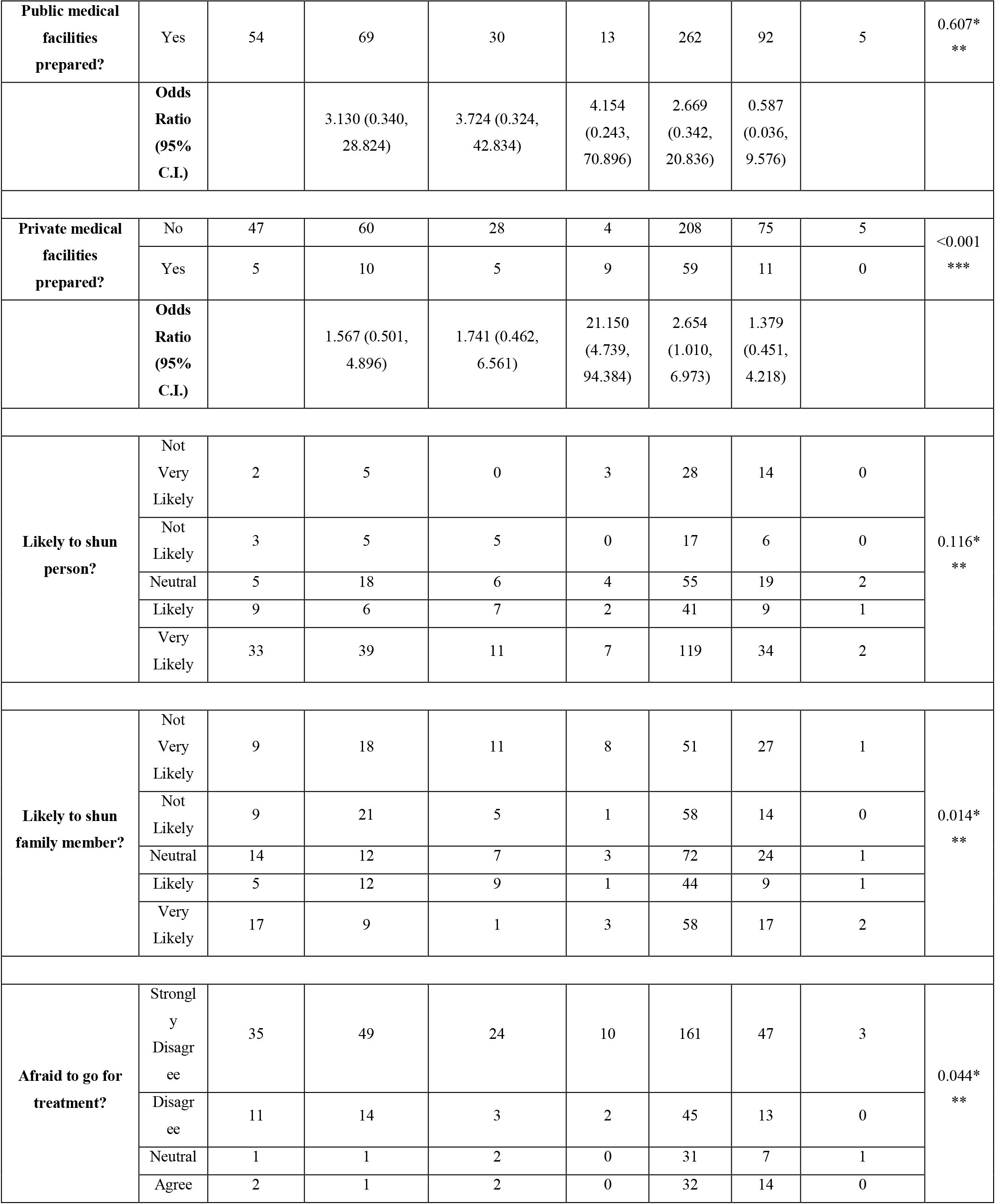
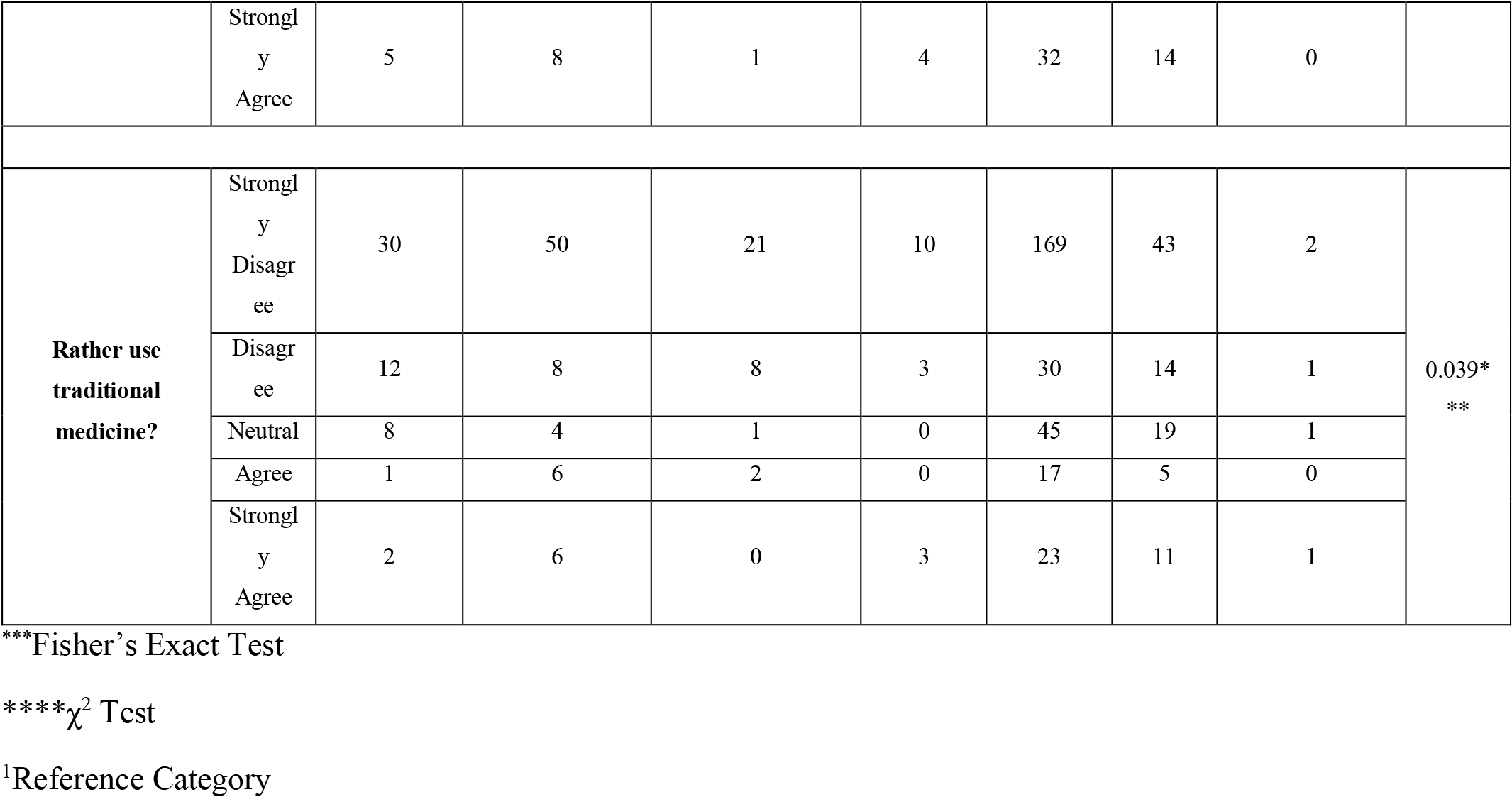
Association of Attitudes toward Ebola and Occupation.

**Table 5.** Association of Attitudes toward Ebola and Education.

| Variable |  | Education |  |  | p-value |
| --- | --- | --- | --- | --- | --- |
|  |  | <sup>1</sup>Primary | Secondary | Tertiary |  |
| **Public medical facilities prepared?** | No | 15 | 223 | 311 | 0.216**** |
|  | Yes | 2 | 11 | 11 |  |
|  | **Odds Ratio (95% C.I.)** |  | 0.370 (0.075, 1.823) | 0.265 (0.054, 1.305) |  |
| **Private medical facilities prepared?** | No | 9 | 179 | 257 | 0.004**** |
|  | Yes | 8 | 48 | 49 |  |
|  | **Odds Ratio**<br>**(95% C.I.)** |  | 0.302 (0.111, 0.824) | 0.214<br>(0.079,<br>0.583) |  |
| **Likely to**<br>**shun**<br>**person?** | Not Very Likely | 3 | 34 | 18 | 0.006*** |
|  | Not Likely | 1 | 16 | 22 |  |
|  | Neutral | 0 | 47 | 64 |  |
|  | Likely | 1 | 33 | 45 |  |
|  | Very Likely | 9 | 89 | 161 |  |
| **Likely to**<br>**shun family**<br>**member?** | Not Very Likely | 2 | 56 | 73 | 0.909*** |
|  | Not Likely | 2 | 42 | 70 |  |
|  | Neutral | 5 | 59 | 73 |  |
|  | Likely | 3 | 37 | 47 |  |
|  | Very Likely | 3 | 45 | 64 |  |
| **Afraid to go**<br>**for**<br>**treatment?** | Strongly Disagree | 4 | 128 | 207 | <0.001*** |
|  | Disagree | 2 | 31 | 59 |  |
|  | Neutral | 3 | 27 | 22 |  |
|  | Agree | 1 | 13 | 18 |  |
|  | Strongly Agree | 5 | 40 | 22 |  |
| **Rather use**<br>**traditional**<br>**medicine?** | Strongly Disagree | 3 | 137 | 197 | 0.013*** |
|  | Disagree | 2 | 31 | 48 |  |
|  | Neutral | 5 | 32 | 46 |  |
|  | Agree | 1 | 19 | 16 |  |
|  | Strongly Agree | 4 | 21 | 19 |  |

\*\*\*Fisher's Exact Test
\*\*\* $\chi^2$ Test
<sup>1</sup>Reference Category

### Preparedness

Further Chi-Squared Tests were carried out to determine the dependence between having a preparedness plan for an outbreak of EVD and demographic factors. In the cases where at least 20% of the cells had expected counts less than 5, Fisher’s Exact Test was used. It was found that preparedness did not depend on demographic factors (p-values > 0.05).

### Obtaining Information about Ebola

Of the 598 respondents who responded to this question, 499 (83.4%) obtained information from various forms of media. Of the 499 responders, the majority (53.3%) were under 30 years of age. However, it was found that the method by which information was obtained was independent of the individual’s age (p-value = 0.673).

## DISCUSSION

The study’s response rate of 67% presented some useful findings about the public’s knowledge and attitudes towards Ebola.

Females outnumbered males but this was somewhat expected since the data collection took place at shopping areas during the day time when more men than women are likely to be at work. The study revealed some useful findings for improving health and healthcare during an outbreak. For example, the significant difference in knowledge scores noted between professional and sales occupations could direct health education measures towards certain occupational groupings. There was no association between age group and knowledge scores; in Nigeria, sub optimal knowledge was found among secondary school students despite a case of imported EVD in 2014 (8).

There was a significant association between age group and being afraid to seek treatment. Fear of seeking treatment suggests a lack of knowledge. The findings show that education level was also significantly associated with being afraid to seek treatment.

Occupational grouping and educational category were associated with likelihood to shun someone with the disease, being afraid to go for treatment and choice of traditional medicine. Education has been found to influence people’s knowledge about EVD in Nigeria (9). In Uganda being educated beyond primary level was a significant predictor of awareness towards EVD and Marburg (10). Our survey showed a significant association between education level and knowledge about EVD. These findings have potentially serious implications for the health system during an outbreak and in peace time as this factor likely applies to all disease and conditions. When patients are prepared to anticipate an untoward situation, they are more likely to cope effectively (11). If persons do not present for treatment, their risk of a poor outcome increases. If they are likely to shun those who are ill, the likelihood of spread would be greater since infection control measures would not be maintained. A knowledge attitude and practices survey undertaken in Liberia (12) found that more than 90% of responders were fearful of EVD patients and of those who live with EVD patients. Choosing to attend traditional healers will lead to further spread as these practitioners would not usually have personal protective equipment and the appropriate training in their usage.

A cause for concern was the finding of the likelihood to shun someone with the disease if you were in a caring occupation. The health workforce density in Guinea, Sierra Leone and Liberia which were the three most affected countries is less than 1.5 per 10 000 population, 2.2 per 10 000 population and less than 3.7 per 10 000 population, respectively (13). In comparison, in TT there are 8 physicians per 10000 people; there are 26 per 10 000 in the USA (14). According to WHO at least 23 physicians, nurses and midwives per 10 000 population are needed for well – functioning primary healthcare interventions (15). The workforce should be well-qualified, capable, responsive and efficient. Lack of a skilled workforce was found to be a major factor in controlling the outbreaks. Lack of knowledge about basic infection prevention and control measures among health care workers was a critical factor in the outbreak response. Lack of PPE and knowing who to report to affected the outcome of the outbreaks (7). Training and rehearsing are critical to a smooth response (16,17,18). However it should be noted that in several studies health workers indicated a reluctance to present to work under emergency conditions because of competing priorities such as family responsibility.

The Government and Public Health practitioners need to change the mindset of the public by providing training and education. When persons are educated and possess the skills to manage a disease which have been rehearsed the fears become less.

Responders obtained their information from various media sources. Media coverage can have a great impact. Governments may therefore use this route for dissemination of clear, accurate information during outbreaks (19).

The little confidence in the health system for handling an outbreak was attributed to a perception that facilities are outdated along with an indifferent attitude. It is noteworthy that responders also had little confidence in private facilities in addition to the public facilities. It seems the Government of TT needs to act to change this perception and /modify their current strategy to inform and educate the public.

The study shows many potential areas for capacity strengthening not only for EVD but for any disease/event that poses a threat to national and thus global health security. The responders are not totally incorrect in their perception that the country is not prepared for an outbreak, since Trinidad and Tobago has not yet fully met its obligations under the IHR 2005.

To build a resilient health system there is a need for community engagement which can aid containment efforts during an outbreak. In addition to the healthcare workers who need to train and to exercise their preparedness plans, the community needs to be involved in the planning and preparedness. In addition to health system strengthening in terms of surveillance and response capacity to early detect and manage outbreaks, the community needs to be engaged and made aware of the risks for each potential threat and of their roles and responsibilities in securing national health security. This could address the issue of staff not presenting to work since community involvement in planning could positively influence decisions healthcare workers take during an emergency with respect to work and surge capacity. In Nigeria respondents suggested sensitisation of communities for effective prevention and control. A survey in the USA stated the need for honest accurate information, ‘*even if that information worried people*.’ The survey report indicated that ‘*These results suggest the value of proactive communications designed to inform the lay public’s decisions, thoughts, and emotions, and informed by concurrent surveys of their responses and needs*’ (20).

### Limitations

The study used convenience sampling which could affect the generalizability of the findings. In terms of the response rate, the non-responders were in the minority. The demographic characteristics (age, sex) of the non-responders were consistent with those of respondents and hence we could have some confidence in the findings from the 622 responders.

### Conclusion

Ebola virus disease was chosen for this study as this disease was of grave concern at the time of the outbreaks since the disease left the African continent for the first time. To quote Dr Keiji Fukuda, a WHO expert, *“what happens in Africa affects your neighborhood, either directly or indirectly…the IHR are founded on that principle…but the world beyond public health remains largely unconvinced of it. To really make strides against diseases like Ebola, we have to convince people that there is no “over there” for infectious pathogens in a globalized world* (2).

We therefore all need to be prepared at all times which means rehearsals and drills in the form of table top exercises and multisectoral simulation exercises. Community support and involvement are key. Sensitisation, communication and training are essential. Unfortunately lessons learned from previous outbreaks are not being used to modify existing services and practices and for each new public health event it seems we are starting from square one. An understanding of the community perceptions about a public health threat is what is needed to be able to develop and target information about the disease and its control (21). This study attempted to achieve this, to aid the Government in its capacity building efforts to increase the country’s resilience during a major public health event.

